# High-dimensional mapping of cognition to the brain using voxel-based morphometry and subcortical shape analysis

**DOI:** 10.1101/479220

**Authors:** Hazel I Zonneveld, Gennady V Roshchupkin, Hieab HH Adams, Boris A Gutman, Aad van der Lugt, Wiro J Niessen, Meike W Vernooij, M Arfan Ikram

## Abstract

**Background:** It is increasingly recognized that the complex functions of human cognition are not accurately represented by arbitrarily-defined anatomical brain regions. Given the considerable functional specialization within such regions, more fine-grained studies of brain structure could capture such localized associations. However, such analyses/studies in a large community-dwelling population are lacking.

**Methods:** In 3,813 stroke-free and non-demented persons from the Rotterdam Study (mean age 69.1 (±8.8) years; 55.8% women) with cognitive assessments and brain MRI, we performed voxel-based morphometry and subcortical shape analysis on global cognition and separate tests that tapped into memory, information processing speed, fine motor speed, and executive function domains.

**Results:** We found that the different cognitive tests significantly associated with grey matter voxels in differential but also overlapping brain regions, primarily in the left hemisphere. Clusters of significantly associated voxels with global cognition were located within multiple anatomic regions: left amygdala, hippocampus, parietal lobule, superior temporal gyrus, insula and posterior temporal lobe. Subcortical shape analysis revealed associations primarily within the head and tail of the caudate nucleus, putamen, ventral part of the thalamus, and nucleus accumbens, more equally distributed among the left and right hemisphere. Within the caudate nucleus both positive (head) as well as negative (tail) associations were observed with global cognition.

**Conclusions:** In a large population-based sample, we mapped cognitive performance to (sub)cortical grey matter using a hypothesis-free approach with high-dimensional neuroimaging. Leveraging the power of our large sample size, we confirmed well-known associations as well as identified novel brain regions related to cognition.

## Introduction

Human cognition comprises a variety of important domains including memory, information processing speed and executive function. Cognitive ability is associated with important health outcomes(Eggermont, et al., 2012; Gale, et al., 2008; Halperin, et al., 2008) and varies between individuals and throughout life(Halperin, et al., 2008). It is determined by both genetic and environmental factors(Deary, et al., 2010; Haworth, et al., 2010), which are reflected in the structure of the brain(Davies, et al., 2015; Kramer, et al., 2004).

Many of the initial links between brain structure and cognition arose from clinical observations of patients with localized brain lesions or following surgical interventions(Newman, et al., 2007; Rorden and Karnath, 2004). Subsequent neuroimaging studies have used these observations in hypothesis-driven approaches to study the neural substrate of human cognitive ability including its various domains(Elderkin-Thompson, et al., 2008; Newman, et al., 2007). These studies have primarily focused on aggregate measures over the entire brain regions e.g., volumetric measures of the prefrontal cortex(Salat, et al., 2004; Tisserand, et al., 2004), thalamus(Van Der Werf, et al., 2001) or hippocampus(Van Petten, 2004). However, it is increasingly recognized that the complex functions of human cognition are not accurately represented by anatomical regions that are arbitrarily defined based on macroscopical landmarks or histological microstructure, e.g. Brodmann areas(Bola and Sabel, 2015; Bressler and Menon, 2010; Park and Friston, 2013). Moreover a considerable functional specialization typically exists within such regions. For example, in Alzheimer’s disease the size of hippocampal subfields contains information important for cognition beyond the gross hippocampal volume(de Flores, et al., 2015; La Joie, et al., 2013; Lindberg, et al., 2012). The thalamus comprises more than 60 cytoarchitectonically and functionally distinct nuclei, all of which have a different pattern of anatomical connections to other brain regions (Fama and Sullivan, 2015; Schmahmann and Pandya, 2008).

An alternative to hypothesis-driven analyses are hypothesis-free approaches that interrogate brain structure at the highest resolution and provide the opportunity to explore the association beyond just aggregated measures. For instance, voxel-based morphometry (VBM) studies volumetric differences at the level of the voxel, the smallest unit of measure of an MRI scan. In recent years, a large body of literature has emerged that employs VBM and related techniques to study how brain structures relate to cognition(Burgaleta, et al., 2014; Fleischman, et al., 2014; Ruscheweyh, et al., 2013; Squarzoni, et al., 2012; Tisserand, et al., 2004). However, still several knowledge gaps remain: First, many hypothesis-free brain imaging studies are performed in relatively small studies, thereby running the risk of false-positive findings or non-significant results. Larger sample sizes can overcome this restriction and yield more robust findings as well as generalize the previous. Second, in addition to VBM analysis, the shape of subcortical structures allows to study the brain regions beyond just volumetric measures(Roshchupkin, et al., 2016b) and may represent underlying subfield organization(Wang, et al., 2008).

Therefore, using hypothesis-free approaches of voxel-based morphometry and shape analysis we performed a fine-mapping of cognitive ability to (sub)cortical grey matter on magnetic resonance imaging (MRI) in a large population-based sample of middle-aged and elderly subjects.

## Materials and methods

### Study population

This study was embedded within the Rotterdam Study, an ongoing prospective population-based cohort designed to investigate chronic diseases in the middle-aged and elderly population(Ikram, et al., 2017). The cohort started in 1990 and comprised 7,983 participants aged ≥55 years. In 2000 and 2006, the study was expanded and at present comprises 14,926 participants aged ≥45 years. Since 2005, brain MRI was implemented into the study protocol(Ikram, et al., 2015). Between 2009 and 2014 4,140 participants underwent brain MRI and cognitive testing. Examinations in this time period were conducted as one project. We excluded participants due to prevalent dementia (n=42), insufficient cognitive screening (n=21), with cortical infarcts (n=103) or clinical stroke (n=161). In total, 3,813 participants were available for analysis. The Rotterdam Study has been approved by the medical ethics committee according to the Population Study Act Rotterdam Study, executed by the Ministry of Health, Welfare and Sports of the Netherlands. A written informed consent was obtained from all participants.

### MRI acquisition

Brain MRI was performed on a 1.5-T MRI scanner (Signa Excite II, General Electric Healthcare, Milwaukee, WI,USA) using an eight-channel head coil. The protocol included T1-weighted sequence (T1), proton density-weighted sequence, and a T2-weighted fluid-attenuated inversion recovery (FLAIR) sequence, as described extensively in detail before(Ikram, et al., 2015).

### Voxel based morphometry

Voxel based morphometry (VBM) was performed according to an optimized VBM protocol(Good, et al., 2001) and as previously described(Roshchupkin, et al., 2016a). Briefly, all T1-weighted images were segmented into supratentorial grey matter, white matter and cerebrospinal fluid using a previously described k-nearest neighbor algorithm, which was trained on six manually labeled atlases(Vrooman, et al., 2007). All grey matter (GM) density maps were non-linearly registered to the standard ICBM MNI152 grey matter template (Montreal Neurological Institute) with a 1×1×1 mm^3^ voxel resolution. A spatial modulation procedure was used to avoid differences in absolute grey matter volume due to the registration, following by smoothing procedure, using a 3mm (FWHM 8mm) isotropic Gaussian kernel.

### Subcortical shapes

The T1-weighted MRI scans were processed using FreeSurfer(Fischl, et al., 2004) (version 5.1) to obtain segmentations and volumetric summaries of the following seven subcortical structures for each hemisphere: nucleus accumbens, amygdala, caudate, hippocampus, pallidum, putamen, and thalamus(Roshchupkin, et al., 2016c). Next, segmentations were processed using a previously described shape analysis pipeline(Gutman, 2015; Roshchupkin, et al., 2016c). Briefly, a mesh model was created for the boundary of each structure. Subcortical shapes were registered using the “Medial Demons” framework, which matches shape curvatures and medial features to a pre-computed template(Gutman, et al., 2013). The templates and mean medial curves were previously constructed and are distributed as part of the ENIGMA-Shape package (http://enigma.usc.edu/ongoing/enigma-shape-analysis/). The resulting meshes for the 14 structures consist of a total of 27,120 vertices(Roshchupkin, et al., 2016c). Two measures were used to quantify shape: the radial distance and the natural logarithm of the Jacobian determinant. The radial distance represents the distance of the vertex from the medial curve of the structure. The Jacobian determinant captures the deformation required to map the subject-specific vertex to a template and indicates shape dilation due to sub-regional volume change(Roshchupkin, et al., 2016c).

### Assessment of cognitive functioning

Cognitive function was assessed with a cognitive test battery comprising Stroop test(Houx, et al., 1993) (word reading, color naming and a reading/color naming interference task (error-adjusted time in seconds)), which tests information processing speed and executive function; 15-Word learning test (15-WLT)(Bleecker, et al., 1988), which taps into immediate and delayed recall to investigate memory; Letter-digit substitution task (LDST)(Prins, et al., 2005) and Word fluency test (WFT, animal categories), both of which test executive function. The three Stroop tests were natural log transformed due to a skewed distribution. To allow for comparison across cognitive tests, we calculated z scores (subtracting the population mean and dividing by the standard deviation) for each cognitive test. The z scores for the Stroop Tests were inverted because higher scores on the Stroop test indicate a poorer performance, whereas higher scores on the other cognitive tests indicate better cognitive performance. In addition, we also investigated global cognition by calculating a compound score (G-factor) using a principal component analysis on the delayed recall score of the 15-WLT, Stroop Interference Test, LDST, and WFT(Hoogendam, et al., 2014). The G-factor explained 57.2% of the variance in cognitive test scores in the population.

### Other measurements

Attained level of education was collected and expressed in years. Prevalent clinical stroke and dementia were assessed based on a protocol as previously described.(Bos, et al., 2014; de Bruijn, et al., 2015)

### Statistical analysis

For VBM and shape analyis, linear regression models were fitted with age, sex, education and cognitive test value as independent variables and voxel or vertex measure as the dependent variable. We corrected for level of education as a measure of cognitive reserve. For both VBM and shapes analysis we computed the significance threshold based on nonparametric statistic test by performing 10,000 random permutations.(Churchill and Doerge, 1994) After collecting the minimum p-value from every test, they were sorted and the 5% quantile was used (α=0.05) to estimate the p-value significant threshold, while controlling the family wise error (FWE). The resulting values were 2.99 x 10^-7^ for VBM and 9.63×10^-6^ for shapes, which were subsequently divided by the number of cognitive tests (n=8) to account for multiple hypothesis tests correction.

## Results

Characteristics of the study population are presented in **Table 1**. Of the 3,813 participants, 55.8% were women and mean age was 69.1 years (ranging from 51.9 to 97.9 years). Correlations between all cognitive test scores stratified by sex are shown in **Supplementary Figure 1**.

**Table 1.** Characteristics of the Study Population.

| **Total population n= 3,813** |  |
| --- | --- |
| Age, years | 69.1 (8.8) |
| Female | 2129 (55.8) |
| Education, years | 12.7 (3.9) |
| *Left hemisphere, cm<sup>3</sup>* |  |
| Nucleus accumbens | 0.55 (0.09) |
| Amygdala | 1.30 (0.21) |
| Caudate nucleus | 3.37 (0.55) |
| Hippocampus | 3.82 (0.63) |
| Pallidum | 1.46 (0.23) |
| Putamen | 4.56 (0.63) |
| Thalamus | 6.15 (0.73) |
| *Right hemisphere, cm<sup>3</sup>* |  |
| Nucleus accumbens | 0.48 (0.09) |
| Amygdala | 1.39 (0.22) |
| Caudate nucleus | 3.48 (0.56) |
| Hippocampus | 3.84 (0.59) |
| Pallidum | 1.39 (0.24) |
| Putamen | 4.40 (0.61) |
| Thalamus | 6.16 (0.73) |

Data presented as mean (standard deviation) for continuous variables and number (percentages) for categorical variables. The following variable had missing data: education (n=52).

### Voxel-based morphometry analysis

In total 4,081 of the grey matter voxels were significant in relation to at least one cognitive tests and/or global cognition after correction for multiple-testing (**Table 2**). These significant voxels were clustered within different brain structures, and almost exclusively within the left hemisphere. The strongest positive associations of better global cognition (G-factor) with grey matter voxels were found in the left amygdala (156 voxels, minimum (min) p-value 4.2×10^-12^), hippocampus (173 voxels, min p-value 9.6×10^-12^), parietal lobule (517 voxels, min p-value 1.2×10^-10^), superior temporal gyrus (313 voxels, min p-value 1.5×10^-10^), insula (142 voxels, min p-value 7.4×10^-10^), posterior temporal lobe left (101 voxels, min p-value 7.7×10^-10^), postcentral gyrus, inferior and middle frontal gyrus, posterior orbital gyrus and right caudate nucleus (all <25 voxels, min p-value all<2.9×10^-8^) (**Figure 1**).

**Table 2.** Association between cognitive tests and grey matter voxels.

| Cognitive Test | Brain Region | Region size (#voxels) | min p-value | # negative voxels | # positive voxels |
| --- | --- | --- | --- | --- | --- |
| **G-factor** | Amygdala left | 2192 | 4.24E-12 | 0 | 156 |
|  | Hippocampus left | 3183 | 9.62E-12 | 0 | 173 |
|  | Parietal lobe left (including supramarginal and angular gyrus) | 65794 | 1.19E-10 | 0 | 517 |
|  | Superior temporal gyrus central part left | 20954 | 1.48E-10 | 0 | 313 |
|  | Insula left | 21930 | 7.43E-10 | 0 | 142 |
|  | Posterior temporal lobe left | 70660 | 7.68E-10 | 0 | 101 |
|  | Inferior frontal gyrus left | 28631 | 9.18E-10 | 0 | 18 |
|  | Middle frontal gyrus left | 80119 | 2.17E-09 | 0 | 15 |
|  | Postcentral gyrus left | 46092 | 2.92E-09 | 0 | 23 |
|  | Posterior orbital gyrus left | 7439 | 4.46E-09 | 0 | 2 |
|  | Caudate nucleus right | 6170 | 2.86E-08 | 0 | 4 |
| **WLT delayed recall** | Hippocampus left | 3183 | 4.84E-10 | 0 | 466 |
|  | Amygdala left | 2192 | 1.13E-09 | 0 | 385 |
|  | Insula left | 21930 | 1.67E-08 | 0 | 10 |
|  | Gyri parahippocampalis et ambiens left | 6823 | 3.45E-08 | 0 | 1 |
| **WLT immediate recall** | Hippocampus left | 3183 | 2.91E-08 | 0 | 6 |
| **WFT** | Superior temporal gyrus, central part left | 20954 | 8.75E-09 | 0 | 103 |
|  | Parietal lobe left (including supramarginal and angular gyrus) | 65794 | 8.81E-09 | 0 | 81 |
| **Stroop interference** | Superior temporal gyrus, central part left | 20954 | 5.29E-11 | 0 | 442 |
|  | Parietal lobe left (including supramarginal and angular gyrus) | 65794 | 5.67E-11 | 0 | 655 |
|  | Postcentral gyrus left | 46092 | 1.86E-10 | 0 | 80 |
|  | Posterior temporal lobe left | 70660 | 2.00E-10 | 0 | 152 |
|  | Insula left | 21930 | 2.52E-10 | 0 | 137 |
|  | Amygdala left | 2192 | 9.21E-10 | 0 | 46 |
|  | Hippocampus left | 3183 | 1.37E-09 | 0 | 24 |
|  | Caudate nucleus left | 6059 | 1.62E-08 | 0 | 21 |
| **LDST** | Insula left | 21930 | 2.62E-08 | 0 | 7 |
|  | Inferior frontal gyrus left | 28631 | 2.72E-08 | 0 | 1 |
| **Total # positive voxels** |  |  |  |  | 4081 |
| **Total # negative voxels** |  |  |  | 0 |  |
| **Total voxels** |  |  |  |  | **4081** |

All associations are adjusted for age, sex, and education. Abbreviations: G-factor global cognition; WLT Word learning test; WFT Word fluency test; LDST Letter-digit substitution task.

**Figure 1.**
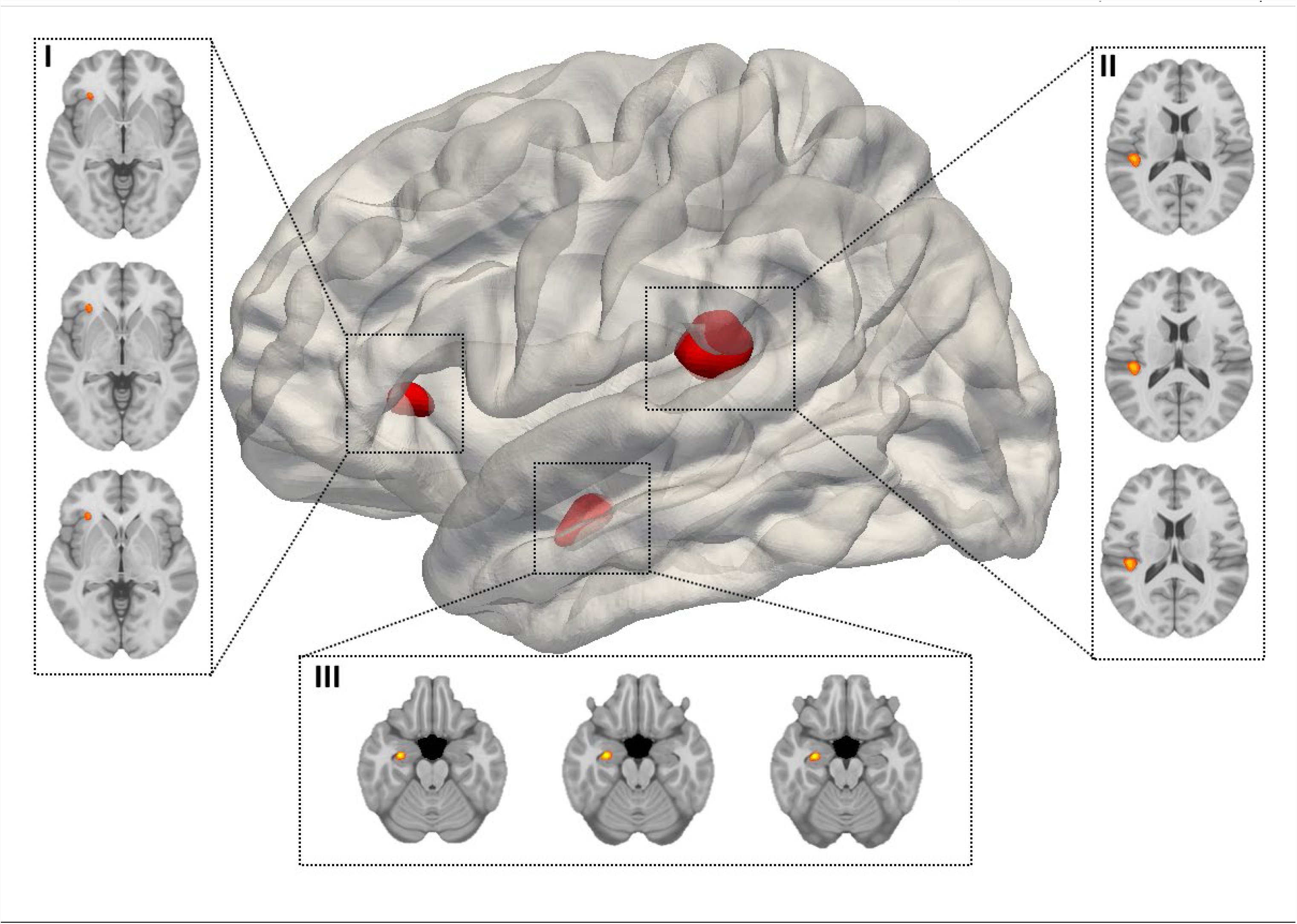
Association of grey matter voxel density with global cognition. Lateral view of the left hemisphere. FWE-significant voxels, indicated by red-yellow, cluster in the insular cortex (I), Wernicke’s area (II), and the hippocampus (III). All associations are adjusted for age, sex, and education. Neurological orientation axial images: left = left.

With respect to the separate cognitive tests, a higher score on the delayed recall task of the 15- WLT was positively associated with grey matter voxels in parts of the left hippocampus (466 voxels, min p-value 4.9×10^-10^), amygdala (385 voxels, min p-value 1.1×10^-9^), insula (10 voxels, min p-value 1.7×10^-8^), and gyri parahippocampalis et ambiens (1 voxel, min p-value 3.5×10^-8^). A higher score on immediate recall score of the 15-WLT was positively related to a small portion of the left hippocampus (6 voxels, min p-value 2.9×10^-8^). We observed that Word-Fluency test (WFT) was associated with superior temporal gyrus (103 voxels, min p-value 8.8×10^-9^), and parietal lobule (81 voxels, min p-value 8.8×10^-9^). We did not observe any association with the Stroop Reading test or the Stroop Color Naming test that survived correction for multiple testing. Stroop interference task harbored significant associations in the left hemisphere including superior temporal gyrus (442 voxels, min p-value 5.3×10^-11^), the parietal lobule (655 voxels, min p-value 5.7×10^-11^), postcentral gyrus (80 voxels, min p-value 1.9×10^-10^), posterior temporal lobe (152 voxels, min p-value 2.0×10^-10^), insula (137 voxels, min p-value 2.5×10^-10^), amygdala (46 voxels, min p-value 9.2×10^-10^), hippocampus and caudate nucleus (both <25 voxels, min p-value 1.6×10^-8^). LDST was associated with a small cluster of voxels in the left insula (7 voxels, min p-value 2.6×10^-8^), and inferior frontal gyrus (1 voxel, min p-value 2.7×10^-8^). All of these results are depicted in **Figure 2A-E**.

**Figure 2.**
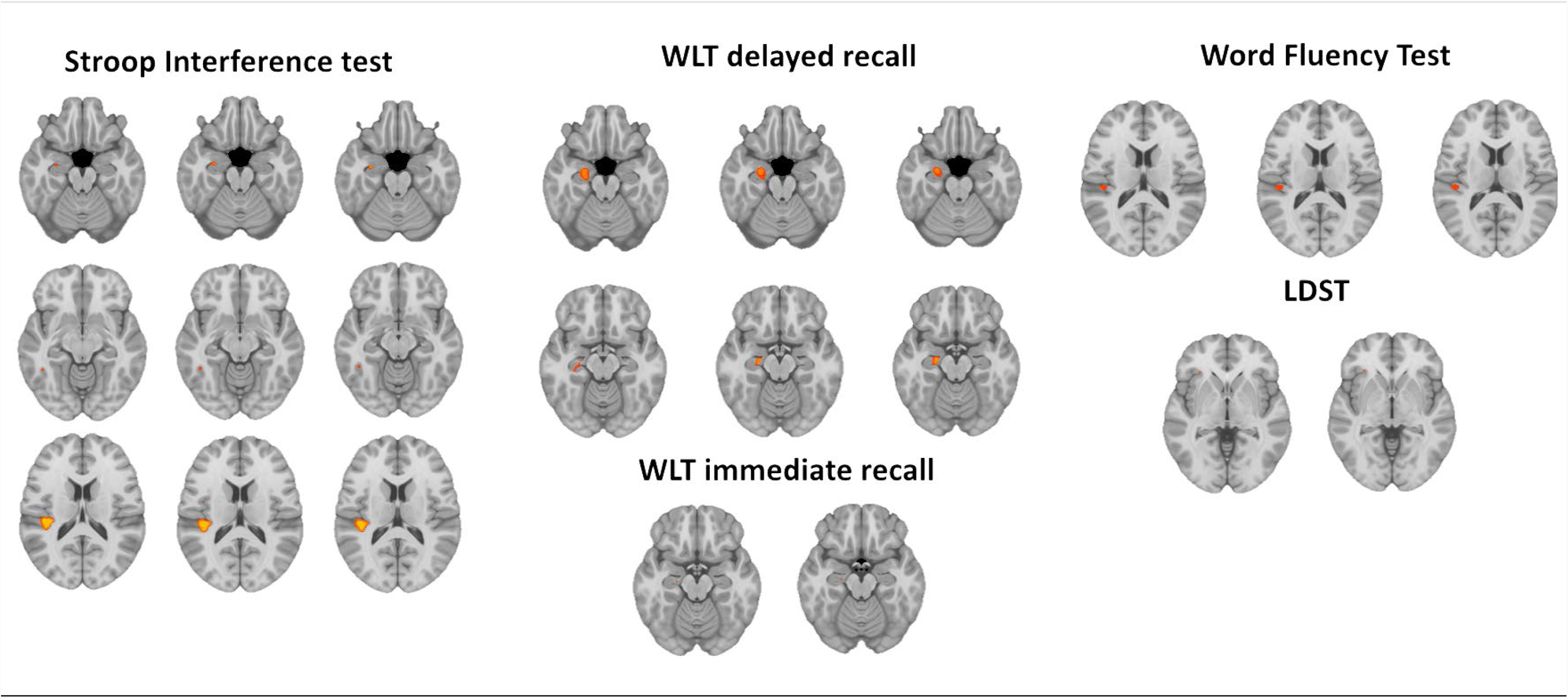
FWE-significant grey matter voxels in relation to cognitive tests. FWE-significant grey matter voxels, indicated by red-yellow, in relation to cognitive tests. Neurological orientation: left ?=left. All associations are adjusted for age, sex, and education. Abbreviations: WLT Word learning test; LDST Letter-digit substitution task.

### Shape analysis

Jacobian determinant and radial distance showed clusters of significant FWE-corrected vertices in relation to cognitive tests, distributed among the left and right hemisphere: 2,819 and 2,298 respectively (**Supplementary Table 2**). The thalamus, caudate nucleus, and putamen harbored most significant associations (**Supplementary Table 2**). Largest significant clusters were found for the Jacobian determinant of the left and right thalamus with Stroop interference task (369 vertices, min p-value 5.8×10^-11^ and 324 vertices, min p-value 6.1×10^-12^ respectively). Global cognition harbored several significant associations, including the Jacobian determinant of the left and right thalamus (281 vertices, min p-value 4.4×10^-12^ and 159 vertices, min p-value 1.9×10^-9^ respectively), and the radial distance of the left and right caudate nucleus (133 vertices, min p-value 2.9×10^-16^ and 78 vertices, min p-value 2.9×10^-13^ respectively) (**Figure 3**). A few inverse associations were observed, primarily between the Jacobian determinant of the caudate nucleus and the WLT (both delayed and immediate recall), and G-factor. Small clusters of vertices (ranging from 1 to 22) were found in the hippocampus with global cognition, WFT and Stroop interference task, but not with the memory tests. **Supplementary Figure 2A-F** shows all significant findings of the shape analysis of subcortical brain structures in relation to the individual cognitive tests.

**Figure 3.**
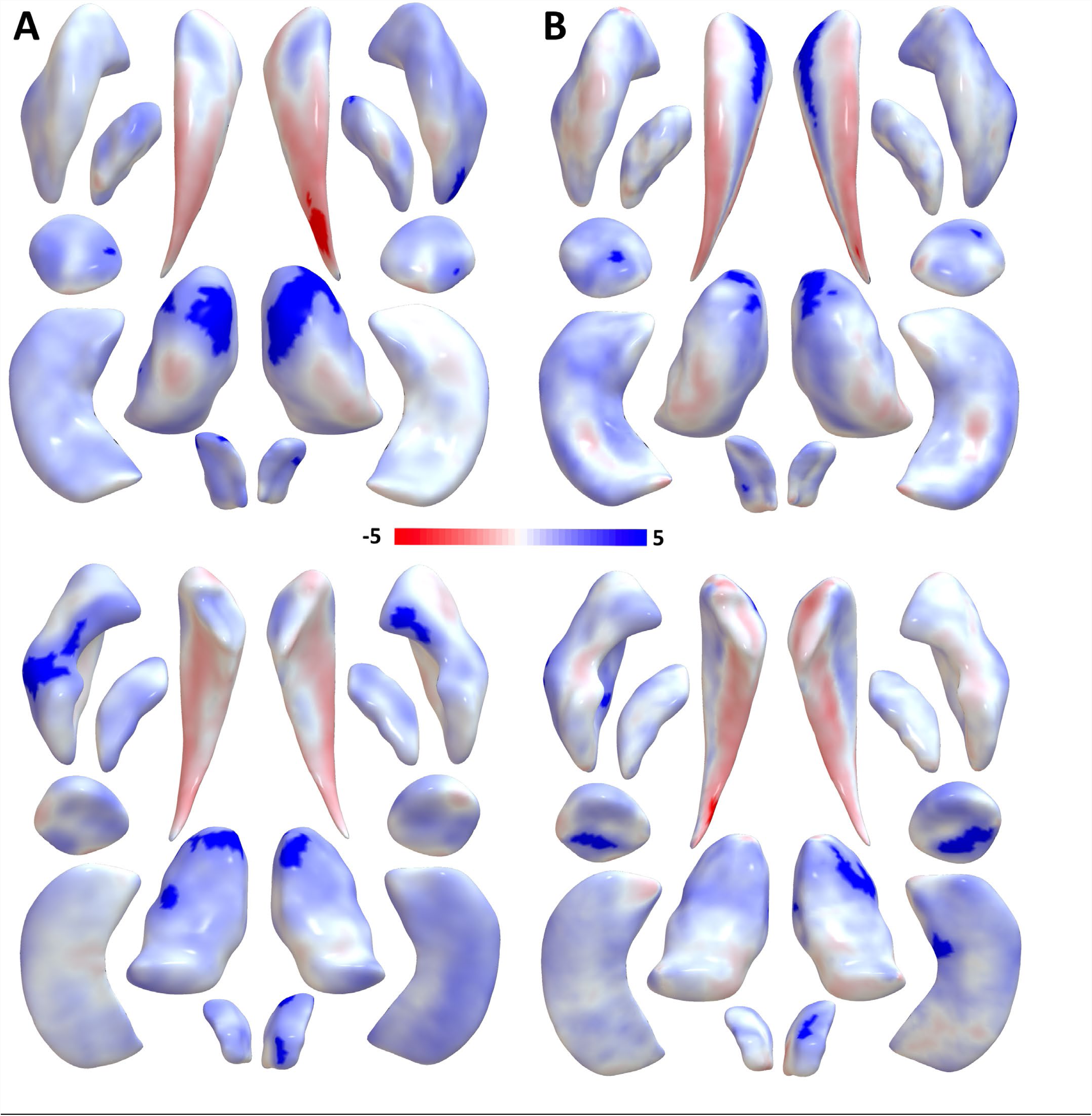
Maps of shape measures of subcortical brain regions in relation to global cognition. Maps show the associations of seven bilateral subcortical structures for the shape measures of Jacobian determinant (Panel A) and radial distance (Panel B), anterior (top row) and posterior (bottom row) view. All associations are adjusted for age, sex, and education. Color map represents the t-statistics and shows the direction of association, with red and blue indicating negative and positive associations respectively. Highlighted regions represent statistically significant vertices

## Discussion

In this large study of community-dwelling adults, we presented the neuroanatomical fine-mapping of seven cognitive tests and global cognition using voxel-based morphometry and subcortical shape analysis. We found that the different cognitive tests significantly associated with grey matter voxels in different brain regions, primarily in the left hemisphere. Moreover, many of the associated regions showed overlap between cognitive tests. Subcortical shape analysis revealed associations primarily within the head and tail of caudate nucleus, putamen, ventral part of the thalamus, and nucleus accumbens, more equally distributed among the left and right hemisphere. Within caudate nucleus both positive (head) as well as negative (tail) associations were observed with global cognition.

Regarding the voxel-based morphometry analysis, we observed three clusters of grey matter voxels to be associated with global cognition. These clusters were found in the left amygdala, hippocampus, parietal lobule, insula, posterior temporal lobe, inferior and middle frontal gyrus, postcentral gyrus and posterior orbital gyrus. Importantly, each of these three clusters was located within multiple anatomic regions. This may emphasize the importance of investigating the association with cognition beyond anatomically defined regions. Global cognition represents the shared variance of the individual cognitive tests, so it is therefore unsurprising that the three significant clusters are also significantly associated with the separate tests. We therefore will discuss our findings in more detail per individual cognitive test below.

Memory research has a long history(Squire and Zola-Morgan, 1988). The medial temporal lobes, and in particular the hippocampus, have long been implicated in episodic memory, with visuospatial memory predominantly associated with the right and verbal memory with the left hippocampus(Burgess, et al., 2002; Frisk and Milner, 1990; Smith and Milner, 1981). In line with this, we found that the 15-Word Learning test, with delayed recall more pronounced than immediate recall, was associated with clusters of grey matter voxels in particular the left hippocampus, as well as in the left amygdala. For decades there has been debate over the question of whether the amygdala is involved in memory(McGaugh, et al., 1996). Task-based resting-state functional MRI studies have shown that the amygdala is considered to play a role in emotional-related memory. However, its role in episodic memory is less known(Phelps, 2004). We did not observe an association between the shape of the hippocampus and cognitive tests measuring memory function. As was shown in a previous study, shape of subcortical structures represents a complimentary phenotype compare to just volumetric measures, with its own genetic architecture (Roshchupkin, et al., 2016c). Therefore the absence of signal may be caused by the fact that the shape of hippocampus has also different functional specialization, which is less sensitive to associations with cognition.

The Stroop interference test, Word Fluency test (WFT) and Letter-Digit Substitution Task (LDST) are all tapping into executive function. The Stroop interference task has been used extensively in studies designed to explore the efficiency of controlled attentional processes(Davidson, et al., 2003). The Stroop effect reflects slowing of response time or increase in error rate when persons are required to respond with the identity if an incongruent stimulus relative to a congruent stimulus(Bugg, et al., 2008). Interestingly, we observed that the Stroop interference task was positively associated with a cluster of grey matter voxels in the left hippocampus. Over the past decades, there has been increasing interest in the contribution of the hippocampus to processes beyond the memory domain(Rubin, et al., 2017). A study in healthy subjects explored the role of the hippocampus for response conflict in the Stroop task by combining intracranial electroencephalography with region of interest-based functional MRI. Researchers found that the hippocampus is recruited during response conflict. Importantly, it remains questionable whether conflict processing can be disentangled from circumstances in which there is conflicting valence or perceptual information, even in experimental studies that thoroughly control for the effect of memory(Ito and Lee, 2016). Moreover, WFT and Stroop interference test showed clusters of significant grey matter voxels in the left hemisphere where the posterior frontal lobe, upper segment of temporal lobe, and parietal lobule (including supramarginal and angular gyrus) intersect. These brain areas are part of Wernicke’s area, a well-known functional language area(Pirmoradi, et al., 2016). The WFT being used in the current study tests semantic fluency. Semantic fluency requires searching for semantic associations within the lexicon(Shao, et al., 2014). Lower scores on semantic fluency tests may therefore also reflect problems with semantic memory, and not only executive function. In line with this, we found WFT to be associated with the left hippocampus, although non-significant after correction for multiple testing.

The LDST and Stroop interference test both showed associations with the left insula, more specifically the dorsal anterior insula. The left insular cortex is involved in consciousness and plays a role in diverse cognitive functions(Dupont, et al., 2003) such as higher cognitive processing and social-emotional processing(Chang, et al., 2013). Anterior insular cortices are among the most commonly activated brain regions across all cognitive tasks(Nelson, et al., 2010). It is also considered to be part of the cognitive control network and it has been hypothesized that this network might form a pathway by which information in the insula, can affect decision making, and therefore influence information processing speed(Cauda, et al., 2011; Deen, et al., 2011).

In line with literature, our shape analysis results indicate that subcortical structures are heterogeneous and consist of functionally diverging sub-regions(de Flores, et al., 2015; Fama and Sullivan, 2015; Lin, et al., 2017; Roshchupkin, et al., 2016c). This is illustrated by, e.g., the caudate nucleus showing that its head and tail differ in their associations with global cognition. It is thought that the head of the caudate nucleus interacts with medial, ventral, and dorsolateral prefrontal cortex as part of the ‘cognitive’ corticostriatal loop, whereas the tail interacts with inferior temporal areas and appears to be involved in visual stimulus processing(Haber, 2016; Lawrence, et al., 1998; Seger, 2013). In addition, our results suggest that the shape of the other subcortical structures are involved in cognition as well, emphasizing the importance of subcortical shape analysis in understanding cognition.

Strengths of our study include the large sample size, the population-based setting and the hypothesis-free approach to be able to fine map cognition to grey matter. Some limitations deserve to be acknowledged. First, because of the cross-sectional design, no conclusions can be drawn regarding the directionality of causality of the associations. Second, our cognitive test battery, limited in time because of the population-based nature, yielded a less extensive cognitive assessment compared to other studies in smaller samples. Third, the current study mainly consists of Caucasians, therefore the generalizability to other ethnicities is limited. Finally, it is well known that several cognitive processes are lateralized to a functionally dominant hemisphere and therefore it would have been interesting to investigate handedness as effect modifier. Unfortunately, in our study we did not have a reliable measure of handedness. In conclusion, in this population-based study of nearly 4000 subjects we mapped cognitive ability to grey matter by using hypothesis-free approaches of voxel-based morphometry and shape analysis. We made the maps of association publicly available (https://neurovault.org/) for any researcher to explore the results or to contrast their findings against. Our results propose that a more fine-grained analysis of brain structure adds to our understanding of cognitive function in normal aging. Future studies are needed to disentangle development and degeneration of the human brain. Additionally, longitudinal assessment of cognitive functioning and grey matter atrophy is needed to study causality.

## Acknowledgements

The Rotterdam Study is funded by Erasmus Medical Center and Erasmus University, Rotterdam, Netherlands Organization for the Health Research and Development (ZonMw), the Research Institute for Diseases in the Elderly (RIDE), the Ministry of Education, Culture and Science, the Ministry for Health, Welfare and Sports, the European Commission (DG XII), and the Municipality of Rotterdam. The authors are grateful to the study participants, the staff from the Rotterdam Study and the participating general practitioners and pharmacists.

## SUPPLEMENTARY MATERIAL

### High-dimensional mapping of cognition to the brain using voxel-based morphometry and subcortical shape analysis

**Content:**

**Supplementary Figure 1:** Correlation between cognitive tests stratified by sex.

**Supplementary Figure 2.** FWE-significant vertices of subcortical brain structures in relation to cognitive tests

**Supplementary Table 1:** Association between cognitive tests and shape measures in the left and right hemisphere.

---

**Supplementary Figure 1.**
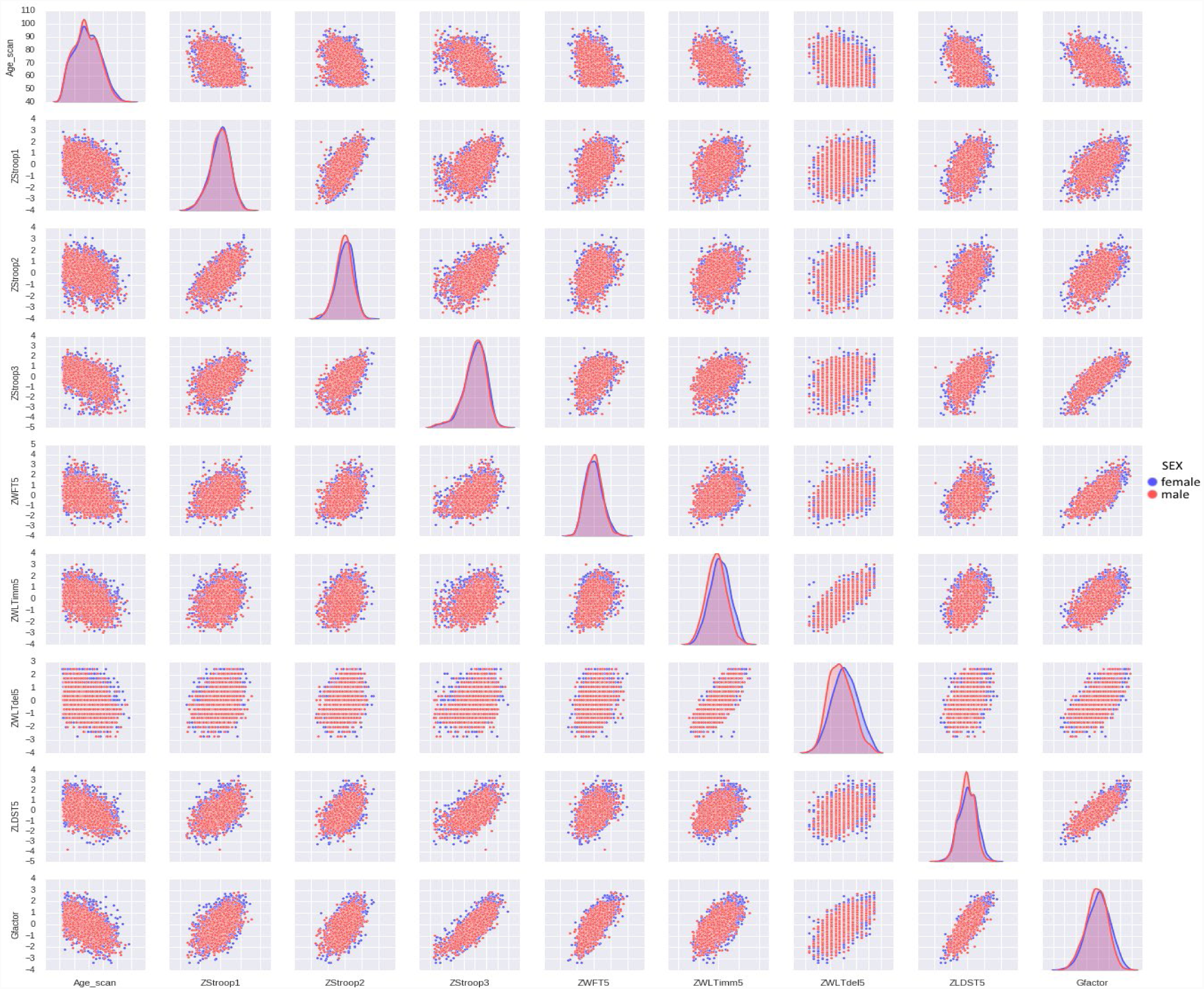
Correlation between cognitive tests stratified by sex.

**Supplementary Figure 2A.**
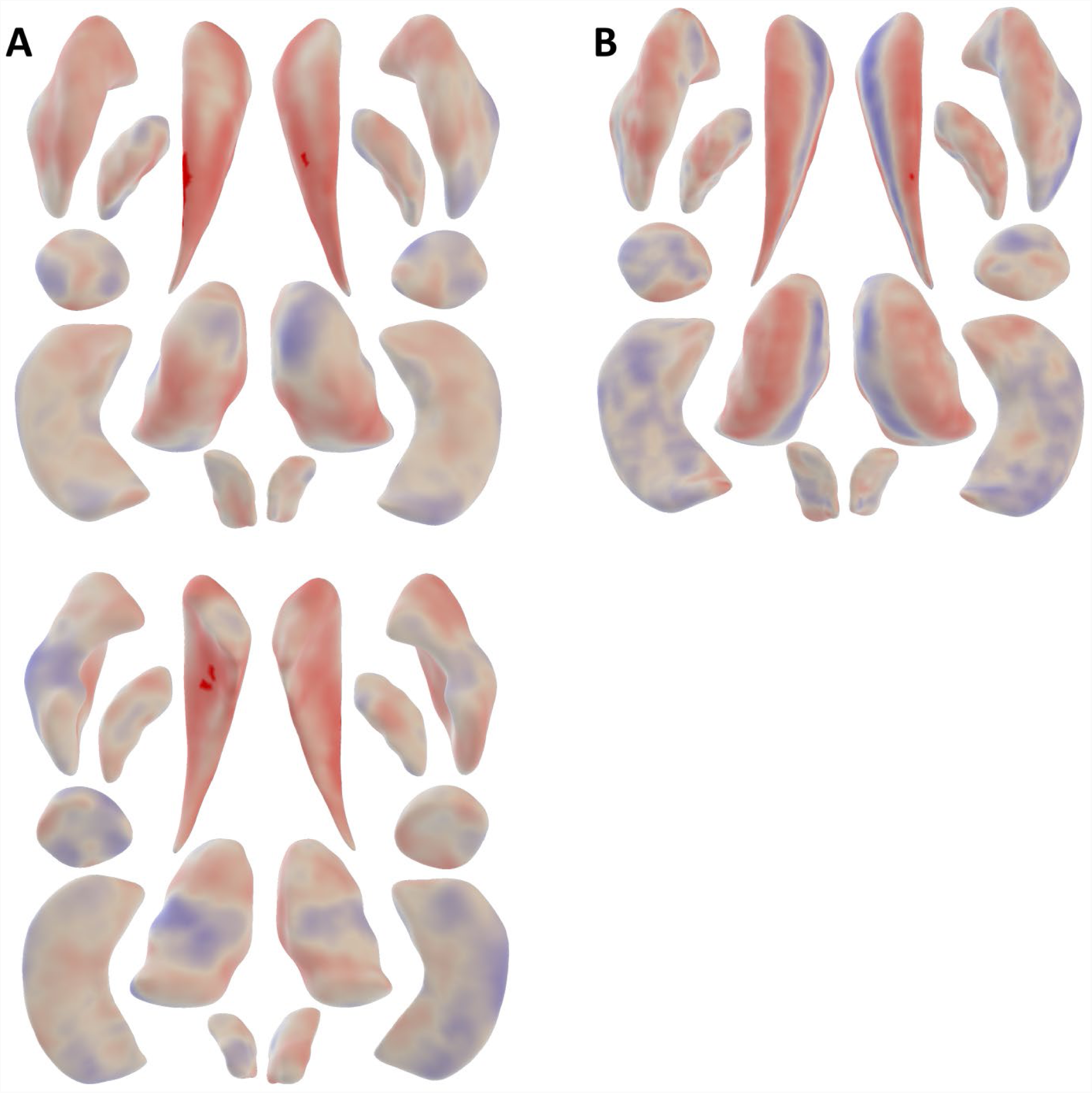
FWE-significant vertices of subcortical brain structures in relation to Word learning test, delayed recall. Maps show the associations of seven bilateral subcortical structures for the shape measures of Jacobian determinant (Panel A) and radial distance (Panel B), anterior (top row) and posterior (bottom row) view. All associations are adjusted for age, sex, and education. Color map represents the t-statistics and shows the direction of association, with red and blue indicating negative and positive associations respectively.

**Supplementary Figure 2B.**
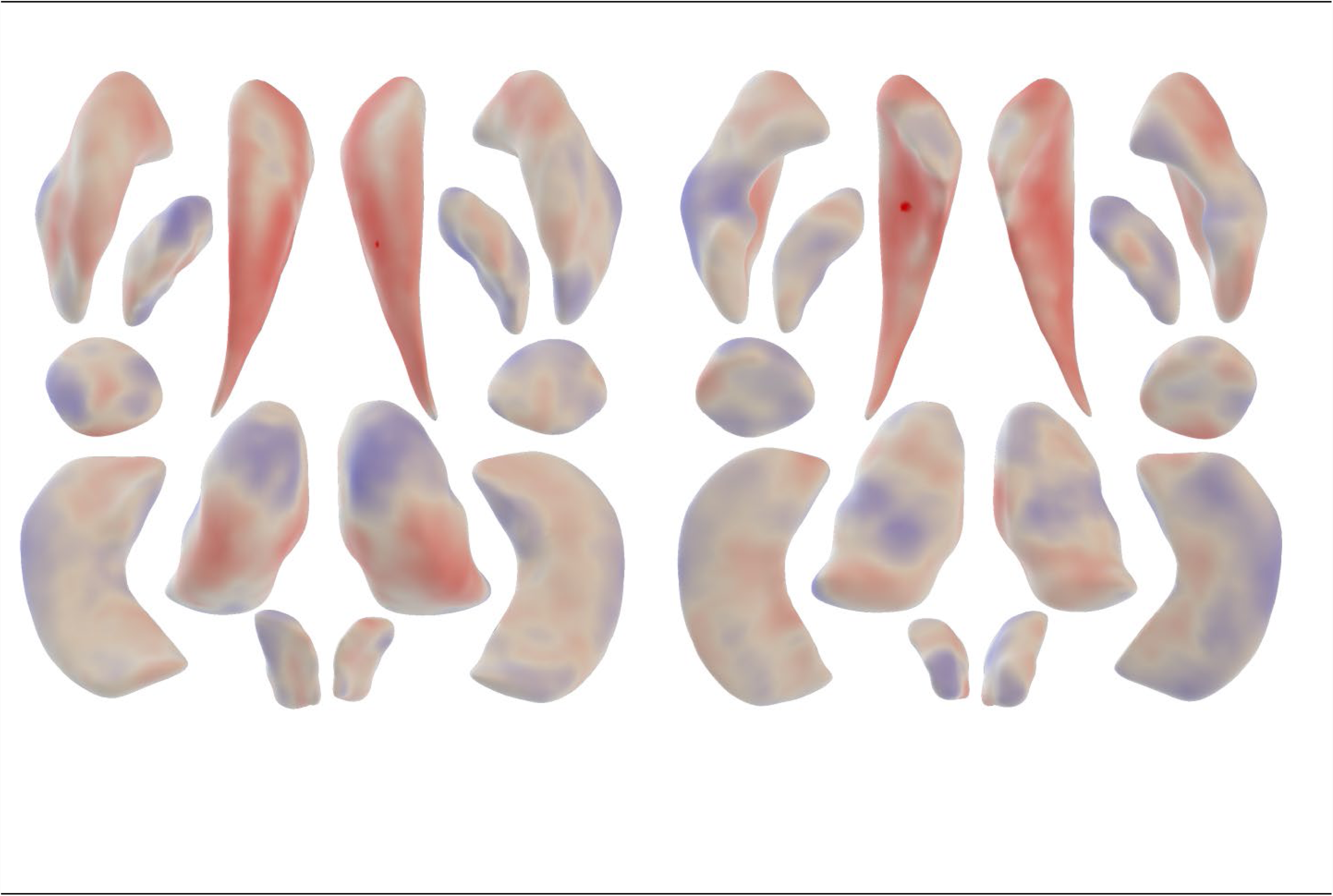
FWE-significant vertices of subcortical brain structures in relation to Word learning test, immediate recall. Maps show the associations of seven bilateral subcortical structures for the shape measures of Jacobian determinant, anterior (left) and posterior (right) view. All associations are adjusted for age, sex, and education. Color map represents the t-statistics and shows the direction of association, with red and blue indicating negative and positive associations respectively.

**Supplementary Figure 2C.**
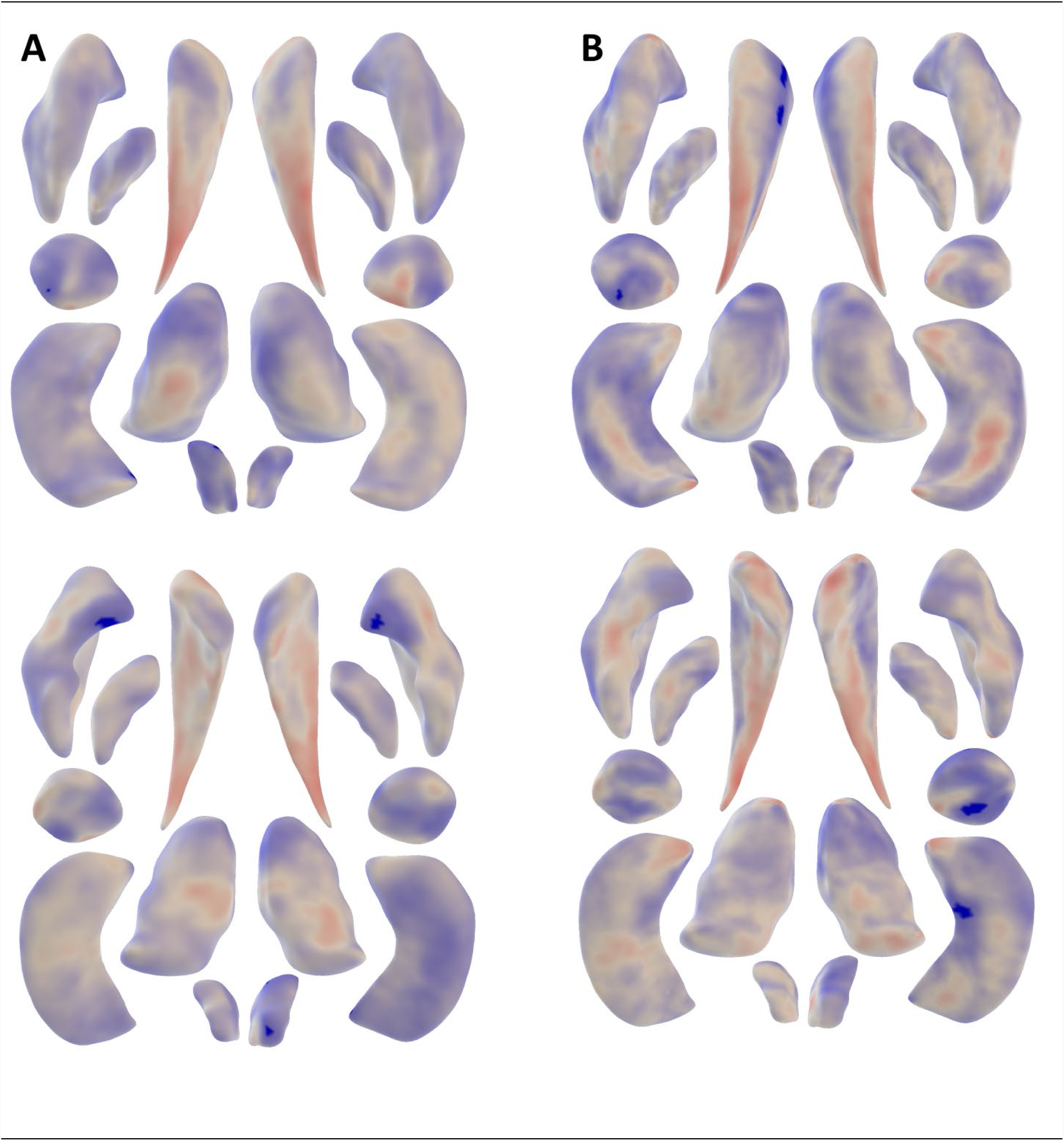
FWE-significant vertices of subcortical brain structures in relation to Word fluency test. Maps show the associations of seven bilateral subcortical structures for the shape measures of Jacobian determinant (Panel A) and radial distance (Panel B), anterior (top row) and posterior (bottom row) view. All associations are adjusted for age, sex, and education. Color map represents the t-statistics and shows the direction of association, with red and blue indicating negative and positive associations respectively.

**Supplementary Figure 2D.**
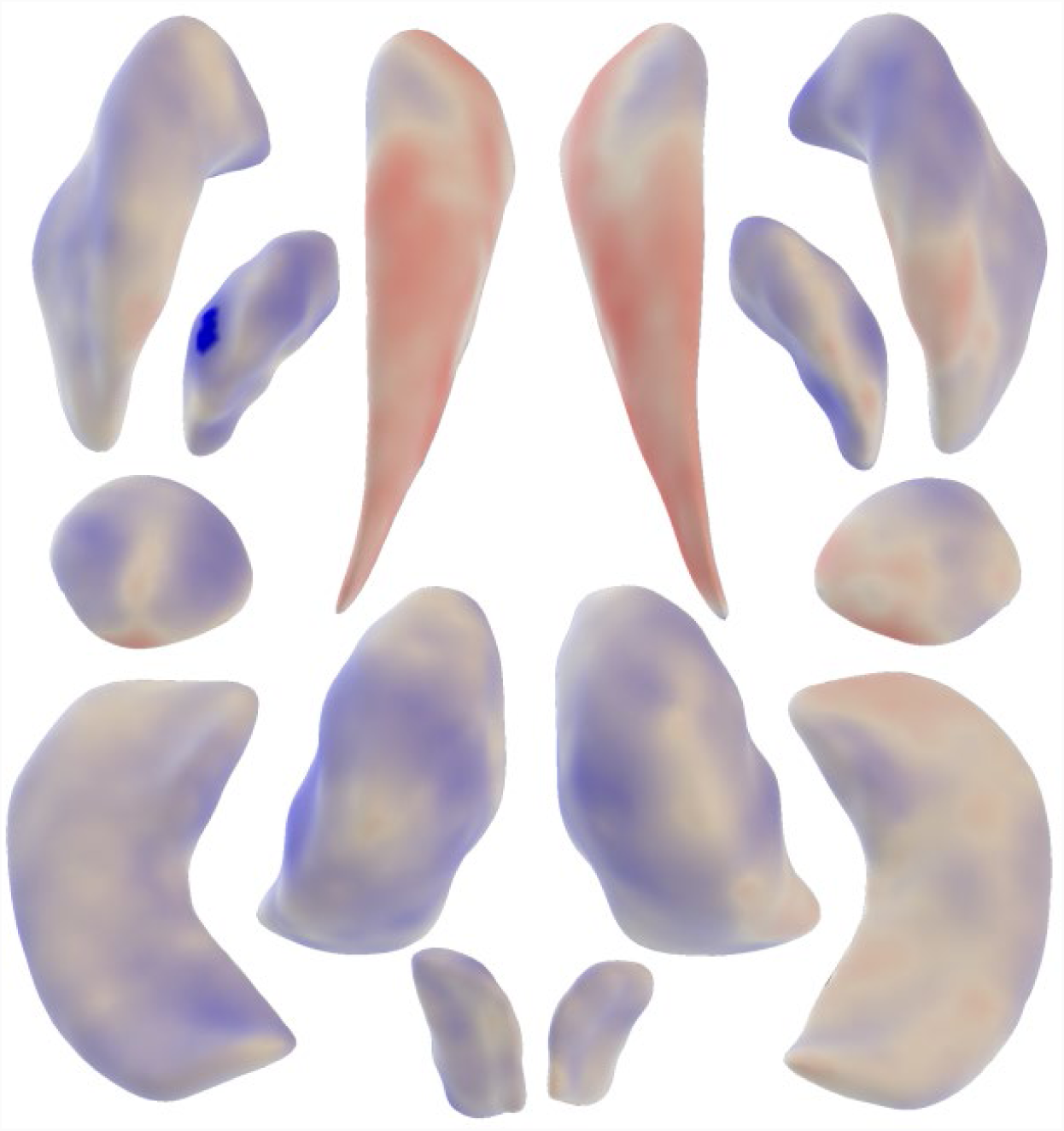
FWE-significant vertices of subcortical brain structures in relation to Stroop color reading test. Maps show the associations of seven bilateral subcortical structures for the shape measures of Jacobian determinant, anterior view. All associations are adjusted for age, sex, and education. Color map represents the t-statistics and shows the direction of association, with red and blue indicating negative and positive associations respectively.

**Supplementary Figure 2E.**
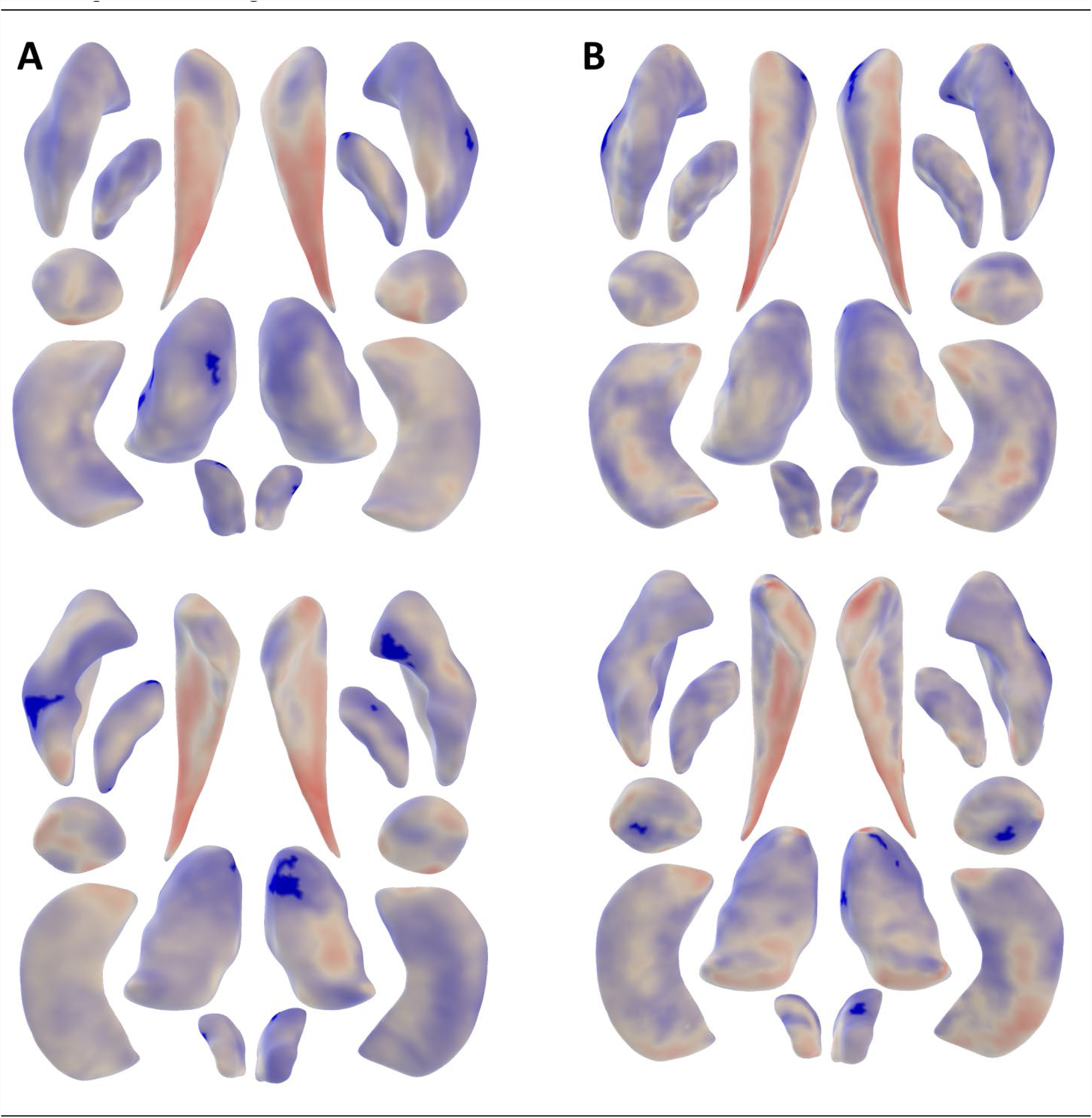
FWE-significant vertices of subcortical brain structures in relation to Stroop color naming test. Maps show the associations of seven bilateral subcortical structures for the shape measures of Jacobian determinant (Panel A) and radial distance (Panel B), anterior (top row) and posterior (bottom row) view. All associations are adjusted for age, sex, and education. Color map represents the t-statistics and shows the direction of association, with red and blue indicating negative and positive associations respectively.

**Supplementary Figure 2F.**
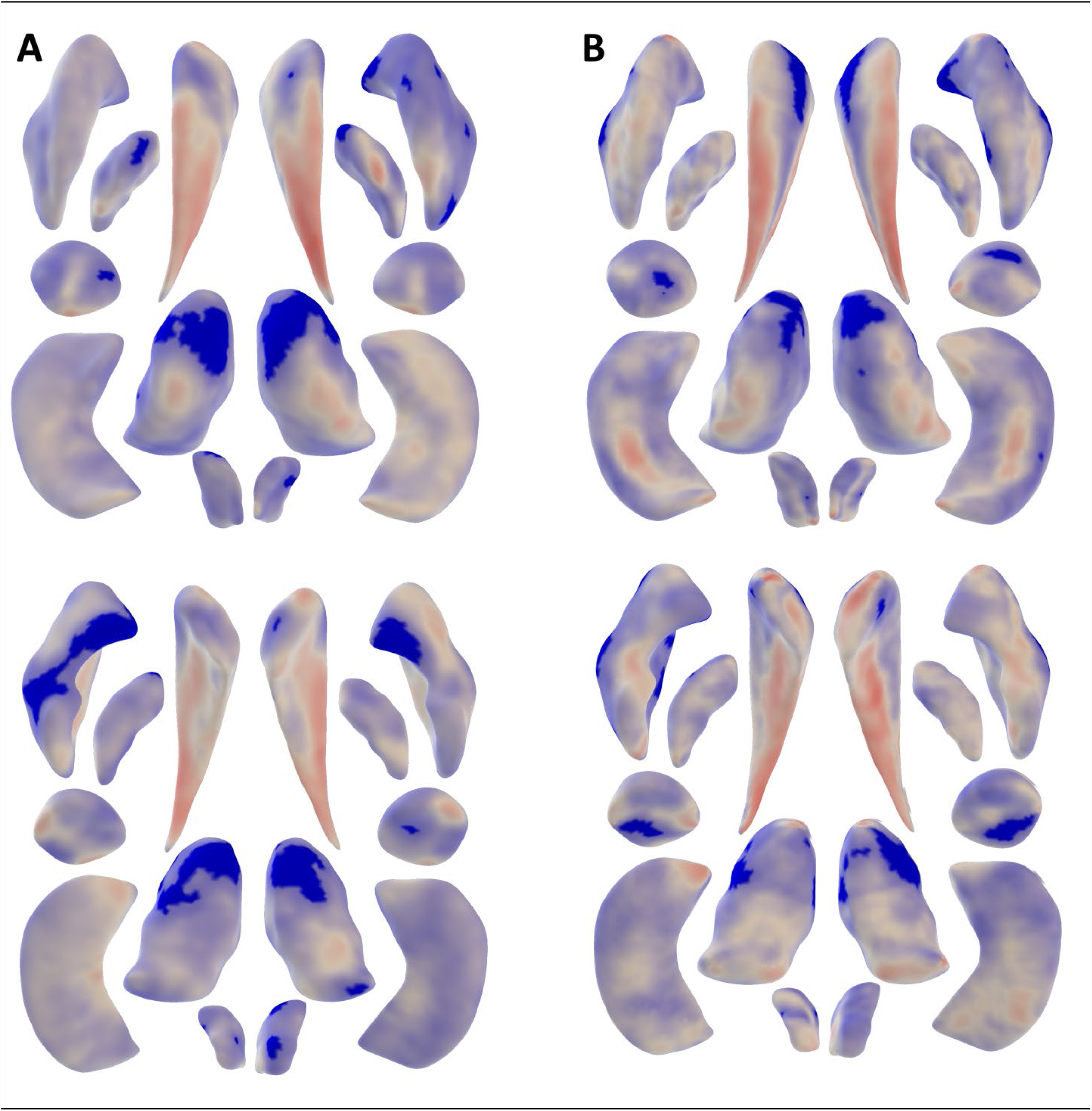
FWE-significant vertices of subcortical brain structures in relation to Stroop interference test. Maps show the associations of seven bilateral subcortical structures for the shape measures of Jacobian determinant (Panel A) and radial distance (Panel B), anterior (top row) and posterior (bottom row) view. All associations are adjusted for age, sex, and education. Color map represents the t-statistics and shows the direction of association, with red and blue indicating negative and positive associations respectively.

**Supplementary Figure 2G.**
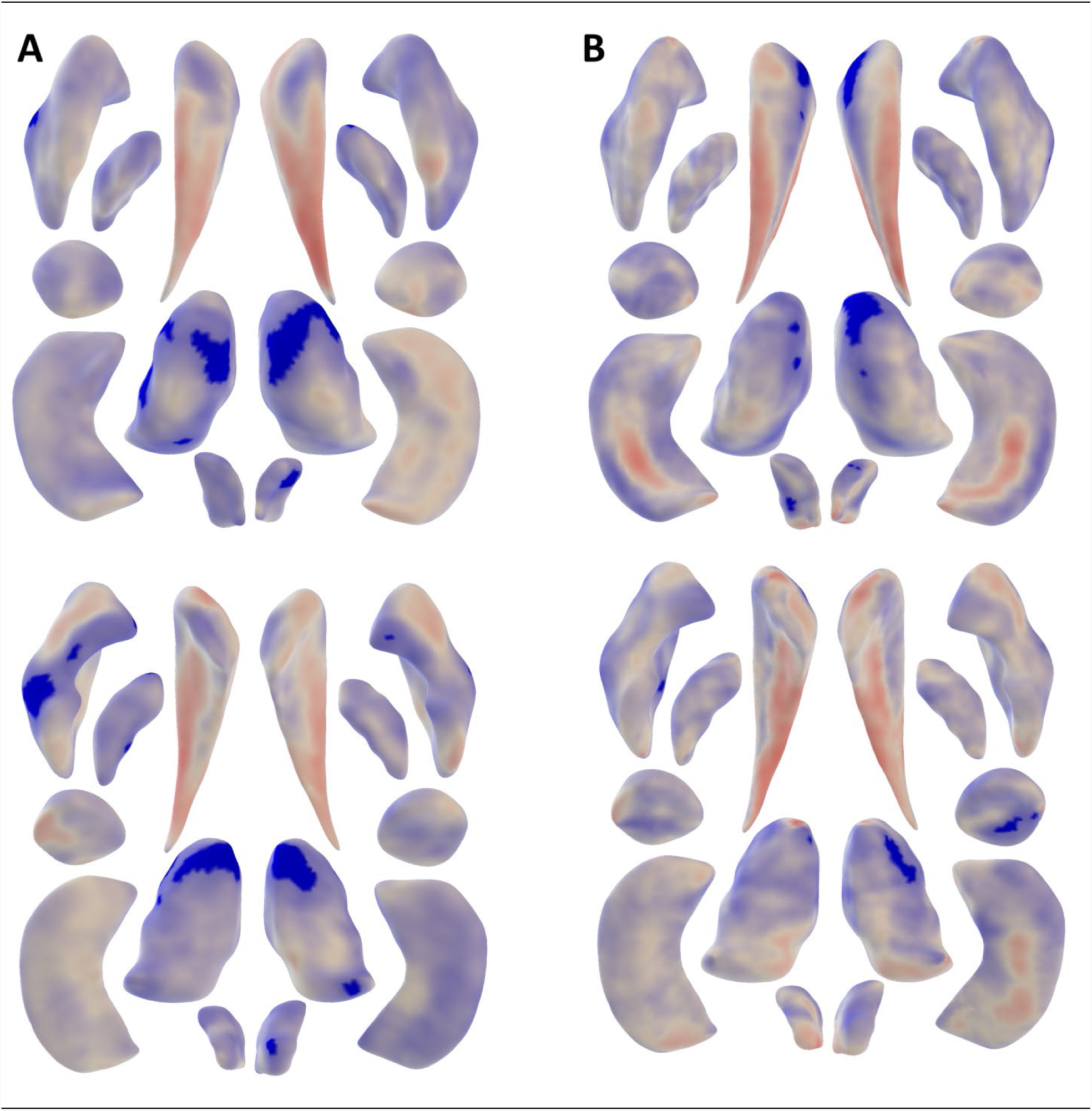
FWE-significant vertices of subcortical brain structures in relation to Letter-digit substitution test. Maps show the associations of seven bilateral subcortical structures for the shape measures of Jacobian determinant (Panel A) and radial distance (Panel B), anterior (top row) and posterior (bottom row) view. All associations are adjusted for age, sex, and education. Color map represents the t-statistics and shows the direction of association, with red and blue indicating negative and positive associations respectively.

**Supplementary Table 1A.**
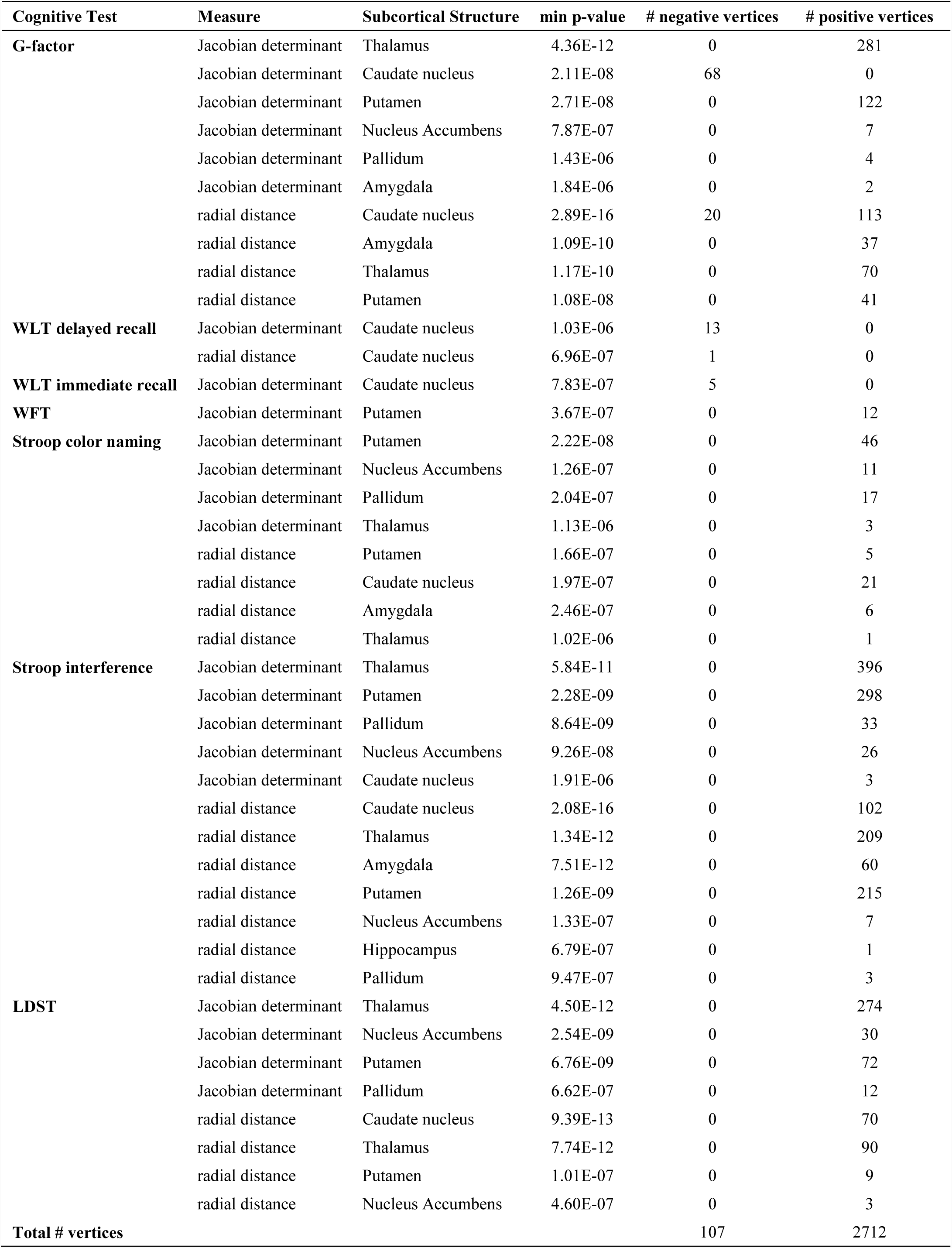
Association between cognitive tests and shape measures in the <u>left</u> hemisphere.

**Supplementary Table 1B.**
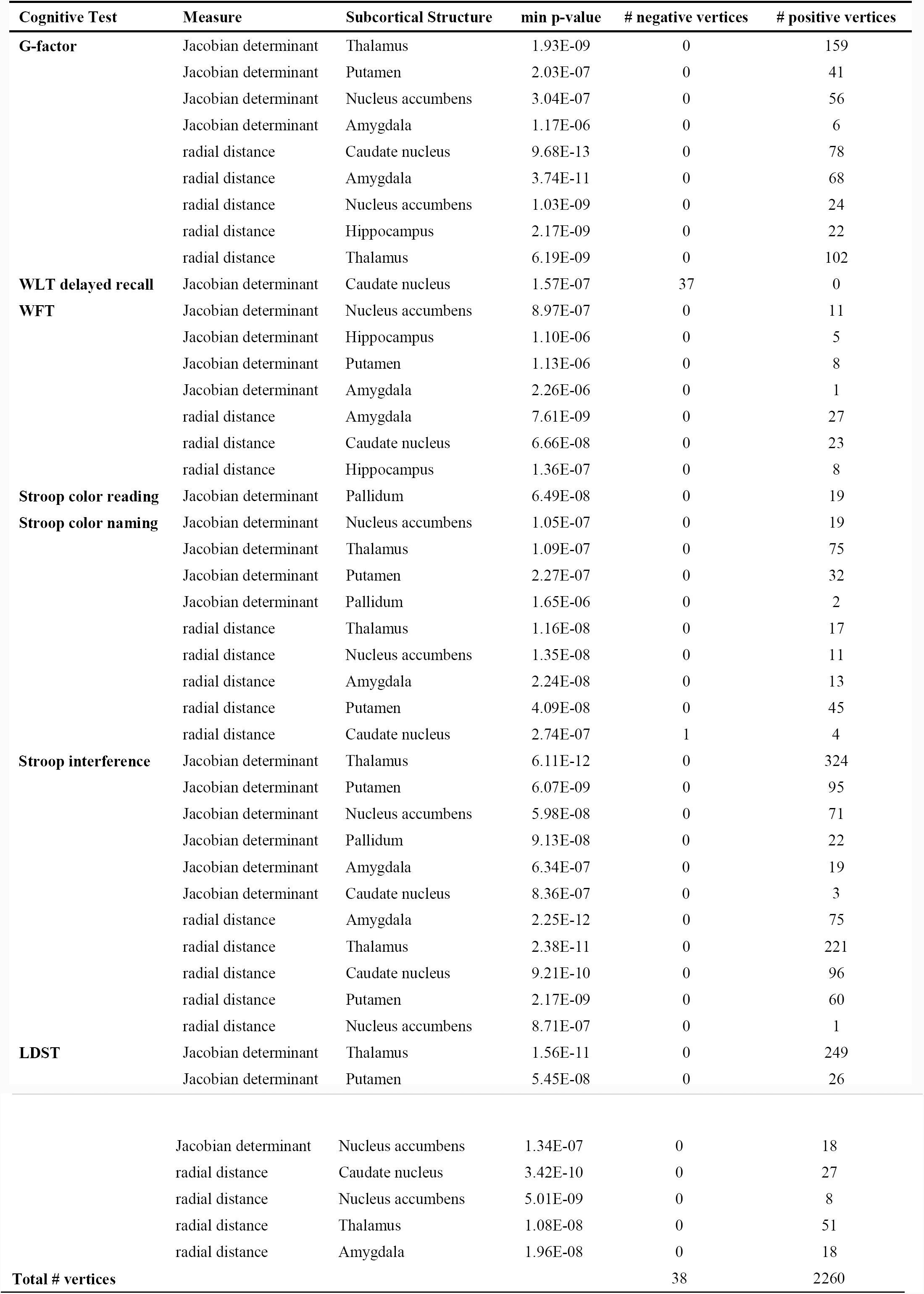
Association between cognitive tests and shape measures in the <u>right</u> hemisphere.

